# Capturing differences in the regulation of LRRK2 dynamics and conformational states by small molecule kinase inhibitors

**DOI:** 10.1101/2022.10.24.513618

**Authors:** Jui-Hung Weng, Wen Ma, Jian Wu, Steve Silletti, J. Andrew McCammon, Susan Taylor

## Abstract

Mutations in the human leucine rich repeat protein kinase-2 (LRRK2) create risk factors for Parkinson’s Disease, and pathological functions of LRRK2 are often correlated with aberrant kinase activity. Past research has focused on developing selective LRRK2 kinase inhibitors. We showed previously that in addition to influencing intrinsic kinase activity, the global conformation of the LRRK2 protein plays a vital role in regulating LRRK2 signaling pathways. Deciphering the allosteric regulation in LRRK2 provides novel strategies for drug discovery. In this study, we combined enhanced sampling simulations with HDX-MS to analyze the inhibitor-induced dynamic changes and the allosteric communications in the C-terminal half of LRRK2, LRRK2^RCKW^. We find that a type I inhibitor (MLi-2) locks the kinase into a closed, active-like configuration, whereas a type II inhibitor (Rebastinib) shifts the kinase to an open, inactive configuration. While both type I and type II inhibitors reduce the kinase activity effectively, they have distinct effects on the LRRK2 conformational dynamics. Specifically, binding of MLi-2 stabilizes the kinase domain in a closed conformation and reduces the global dynamics of LRRK2^RCKW^, leading to a more compact LRRK2^RCKW^ structure. In contrast, binding of Rebastinib stabilizes an open conformation where communication between the N- and C-lobe is severed, which promotes a more extended LRRK2^RCKW^ structure. Rebastinib, based on HDX-MS, creates a more dynamic kinase domain especially at domain interfaces associated with the C-lobe. Our results also reveal the importance of the Dk-helix, which plays a crucial role in propagating communication between the kinase domain and the GTPase domain.

## Introduction

LRRK2 (leucine rich repeat protein kinase-2) is a large 2527 residue multi-domain protein that contains armadillo (ARM), ankyrin (ANK), and leucine-rich (LRR) repeats followed by a tandem Roco type GTPase consisting of a ROC and COR domain, a Ser/Thr kinase (KIN) domain and a C-terminal WD40 domain (Fig 1A). Mutations in LRRK2 cause it to become a risk factor for Parkinson’s Disease (PD), and the pathological functions of LRRK2 correlate mainly with aberrant kinase activity [1-6]. The modulation of LRRK2 kinase activity via the design of small-molecule inhibitors has thus been a central focus for treating PD [7-10], and most of the studies so far have focused on kinase inhibitors. There are two widely studied classes of kinase inhibitors, ATP-competitive Type-I inhibitors that bind to the kinase domain and lock it into a closed and active-like conformation, and Type-II inhibitors that are non-competitive with ATP and typically maintain the kinase in an open inactive conformation [11]. To date, most drug research focuses on type I inhibitors because of their better selectivity [12-14], and many experiments have been conducted to increase their efficacy. However, some potent type I kinase inhibitors, such as MLi-2 (Merck LRRK2 Inhibitor 2), appear to stabilize a disease-like cellular phenotype where LRRK2 accumulates on microtubules [15]. PD mutations such as I2020T with unchanged or reduced catalytic activity dock spontaneously onto microtubules, whereas wild type and the hyperactive mutant, G2019S, only dock in the presence of MLi-2 [16-19]. In contrast, some high affinity type II kinase inhibitors, such as Rebastinib, Ponatinib, and GZD-824, appear to lock LRRK2 into a conformation that is unable to bind to microtubules [20,21]. Both type I and type II LRRK2 inhibitors inhibit LRRK2-mediated phosphorylation of Rab proteins, and both can also stimulate mitophagy, which is negatively regulated by LRRK2 [22]. Only type I inhibitors, however, reduce the phosphorylation of well-studied LRRK2 biomarker sites at the N-terminal region of LRRK2 by inducing dephosphorylation while type II inhibitors do not [23]. This suggests that, in addition to the intrinsic kinase activity towards Rab substrates, the conformation of the LRRK2 protein, likely regulated by the opening and closing of the kinase domain and modulated by binding of 14-3-3 proteins [24,25], plays an important role in mediating the steady state phosphorylation of the biomarker sites, which regulate LRRK2 function [23].

**Fig 1.**
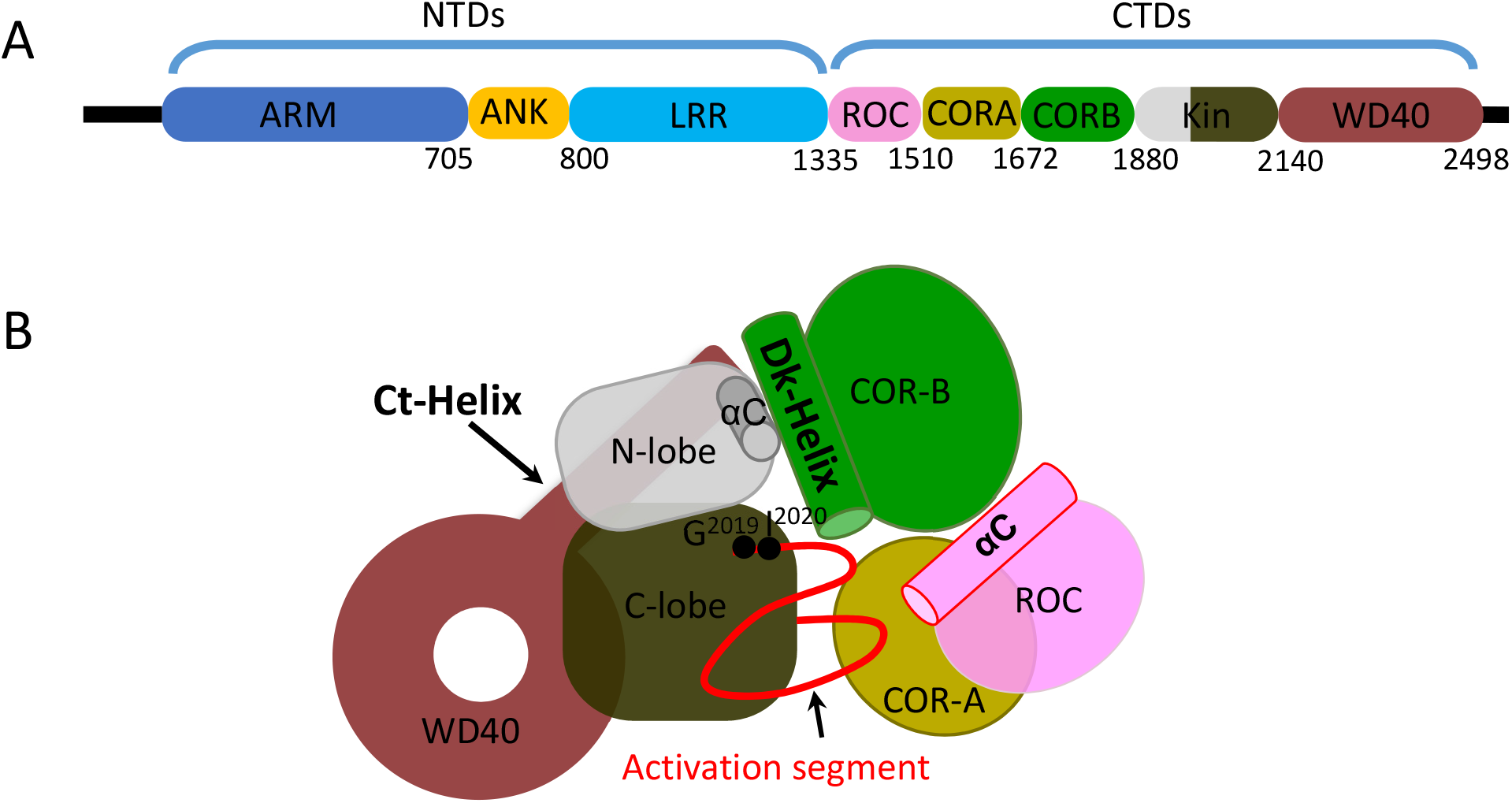
Schematic domain organization of LRRK2. (A) Full length LRRK2 consists of the Armadillo domain (ARM); Ankryn repeat (ANK); Leucine-rich Repeat (LRR); Ras of Complex (ROC), GTPase domain; C-terminal of Roc domain (COR); Kinase domain and WD40 domain. The N-terminal domains (NTDs) contain the ARM, ANK and LRR domains and the C-terminal domains (CTDs) contain the ROC, COR, Kinase and WD40 domains. (B) The model of LRRK2 the CTDs from the Cryo-EM structure shows how the kinase domain is surrounded by the flanking domains.

To date, several high-resolution LRRK2 structures are available of both full length LRRK2 and a truncated version of LRRK2 that contains only the C-terminal domains (LRRK2^RCKW^), and these provide an incredibly valuable resource for delving more deeply into the mechanistic features that regulate LRRK2 structure and function [20,26]. The kinase domain is surrounded by the CORA, CORB, and WD40 domains (Fig 1B), and LRRK2 is one of the only kinases that has a GTPase domain embedded in the same polypeptide. With LRRK2^RCKW^ we can thus capture the direct cross talk between these two major signaling motifs that control so much of biology. Both active and inactive LRRK2 structures have been captured by cryo-EM, and these structures reveal distinct domain movements that resemble the closed and open states of the breathing dynamics [27]. However, no structure of the LRRK2 kinase domain in complex with small-molecule inhibitors has been reported. Other studies on the binding mode between LRRK2 and kinase inhibitors have been mostly based on molecular docking calculations or structures from homolog models of the LRRK2 kinase domain or LRRK2-like mutation studies, and they all focused on the kinase domain alone [28]. We had previously shown that the conformation of the kinase domain plays a crucial role in the intrinsic regulatory processes that mediate subcellular location and activation of LRRK2 [29]. In addition, we identified two helices, the Dk-helix in CORB that docks onto the αC helix of the kinase domain, and the C-terminal helix that spans across the kinase domain, that both shield the kinase domain and regulate LRRK2 conformation and function [30].

In this study, using Gaussian Accelerated Molecular Dynamics MD (GaMD) simulations coupled with Hydrogen Deuterium Exchange Mass Spectrometry (HDX-MS), we studied the C-terminal domains of LRRK2, which include the ROC, COR, Kinase, and WD40 domains (LRRK2^RCKW^), to show how LRRK2 dynamics is affected differently by binding to type I vs. type II kinase inhibitors. These studies show how the N- and C-Lobes function as independent rigid bodies that are stabilized by MLi-2 in a closed and active-like conformation but stabilized by Rebastinib in an open conformation where the two domains are uncoupled. The critical regulatory triad (K1906 in β3, E1920 in the αC helix, and D2017 in the DYG motif) formed in the MLi-2 structure allows the regulatory spine (R-spine) to assemble in an active-like conformation. In contrast, the triad is broken in the Rebastinib-bound LRRK2^RCKW^ structure where D2017 is far from the K1906-E1920 ion pair, and in this structure the R-spine is broken. Our studies also validate the importance of the dynamic features of the Dk-helix, which plays a crucial role in bridging the two catalytic domains, the kinase and GTPase domains. The HDX-MS data revealed that, unlike the type I inhibitor, MLi-2, which reduces the deuterium uptake of the entire kinase domain and the flanking domains that lie in close proximity to the kinase domain, the type II inhibitor, Rebastinib, reduces the local deuterium uptake in the N-lobe but actually increases deuterium uptake in the C-Lobe of the kinase domain. Binding of rebastinib also stabilizes the interfaces among ROC, CORA and CORB domains, while binding to MLi-2 does not. Using GaMD simulations we then validated that the type I inhibitor stabilizes the closed, active-like conformation of the kinase domain and promotes the compact domain orientation of LRRK2^RCKW^. In contrast, the type II inhibitor locks the kinase domain into an open conformation by separating the N- and C-lobes, which in turn stabilizes the domains of LRRK2^RCKW^ in an extended conformation. The dynamic changes in the kinase domain that propagate through the Dk-helix also lead to different conformations of the ROC domain, which potentially affect the GTPase activity. The dynamic and conformational changes described here may also participate in mediating the scaffold and/or oligomerization properties of LRRK2 in signaling, which leads to different LRRK2 function.

## Results

### HDX-MS analysis shows different effects of Type I and Type II kinase inhibitors on LRRK2

To gain insight into how type I and type II inhibitors affect the solvent accessibility of the catalytic domains of LRRK2, we mapped the HDX-MS data onto LRRK2^RCKW^ to compare changes in deuterium uptake in the presence of MLi-2 and Rebastinib (Fig 2). We found that while MLi-2 reduces the deuterium uptake of the kinase domain, Rebastinib shows a different effect on the N-l and C-lobes of the kinase domain. For peptides that cover the CORB-kinase linker (aa 1876-1883), β3-αC linker (aa 1905-1916) and N-Lobe-C-Lobe linker (aa 1948-1958), binding of Rebastinib reduces the H-D exchange but not as effectively as MLi-2. For example, in the linker peptide (aa 1948-1958), Rebastinib reduces its deuterium uptake from 50% to 35%, while MLi-2 reduces the uptake to 25%. In contrast, for the Glycine-rich loop (aa 1884-1893), which is highly protected by MLi-2, the deuterium uptake was unchanged when binding to Rebastinib. For the αC-β4 loop, the low deuterium uptake indicated that it is almost completely shielded from solvent in the WT condition, and both MLi-2 and Rebastinib reduce their deuterium uptake even more.

**Fig 2.**
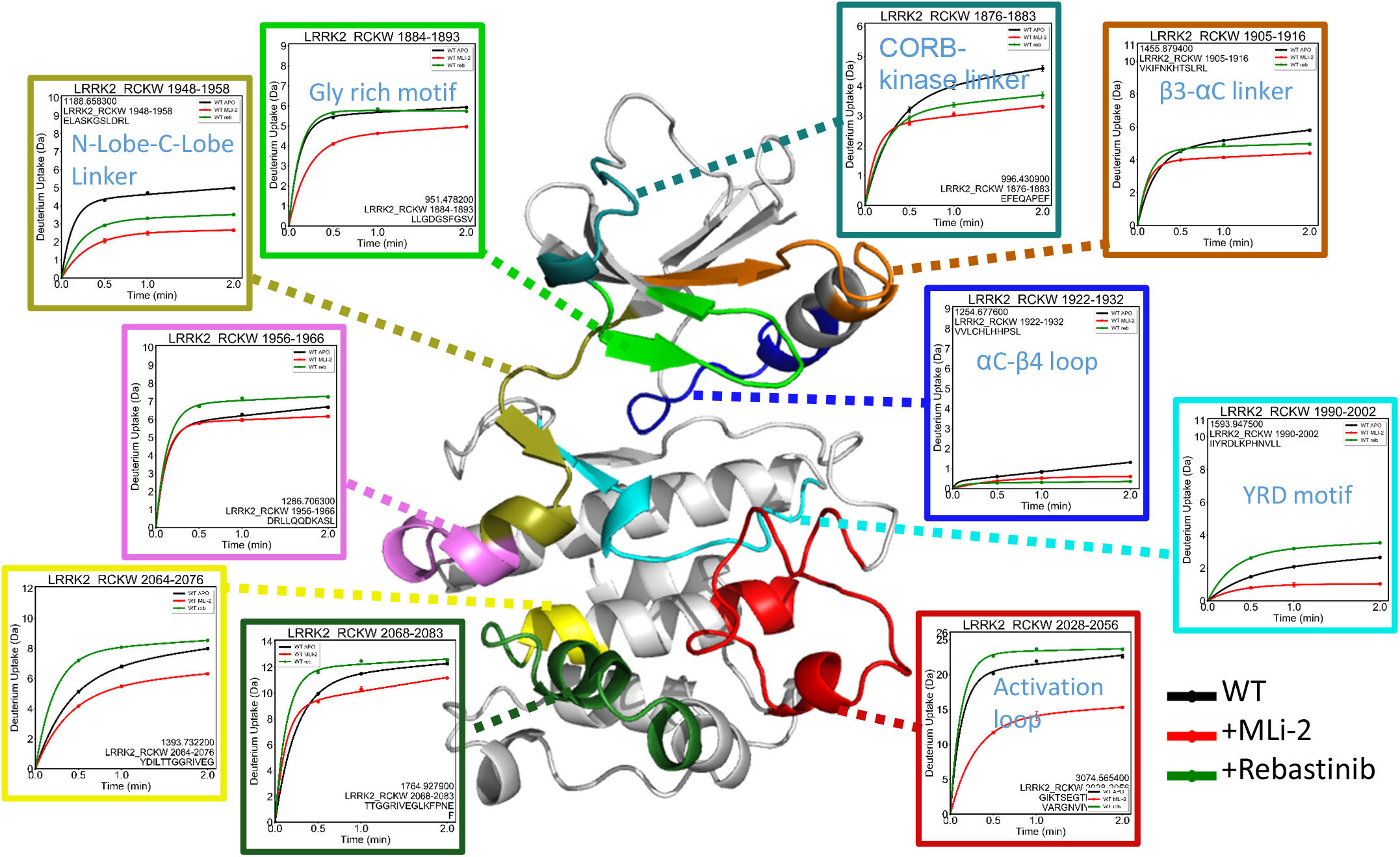
The deuterium uptake of LRRK2 kinase domain. The deuterium uptakes of the selected peptides are plotted and mapped on the kinase model. For all peptides, the uptake is reduced in the presence of MLi-2. Binding of Rebastinib reduces the uptake of CORB-kinase linker, β3-αC linker, N-Lobe-C-Lobe Linker, but the effect is less than that of MLi-2. In contrast, the uptake increases for peptides that cover the C-lobe of the kinase when binding to the Rebastinib.

Interestingly, binding of Rebastinib does not reduce the deuterium uptake in the C-lobe of the kinase domain. Instead, the deuterium uptake is actually increased in several regions in the C-lobe (Fig 2). The activation loop is highly flexible and has high deuterium uptake. The peptide that covered this region (aa 2028-2056) has a higher deuterium uptake when binding to Rebastinib, in contrast to the effect of MLi-2, which has a significant reduction in deuterium uptake. The catalytic loop (aa 1990-2002), including the YRD motif, also becomes more solvent exposed (20% to 35%) when binding to Rebastinib whereas binding of MLi-2 reduced its deuterium uptake to less than 10%. Peptides at the C-terminal end of αF (aa 2064-2076), the αF-αG loop (aa 2068-2083), and the loop between the hinge region and αE (aa 1956-1966) all become more solvent exposed when binding to Rebastinib. Unlike MLi-2 that reduces the deuterium uptake of the entire kinase domain, binding of Rebastinib only protects a localized portion of the N-lobe of the kinase domain from solvent while the C-lobe becomes globally more solvent exposed when binding to Rebastinib.

### Binding of kinase inhibitors changes the conformation of LRRK2

To explore the allosteric impact of inhibitor binding, we carried out three GaMD simulations: LRRK2^RCKW^ (RCKW), LRRK2^RCKW^ in complex with MLi-2 (RCKW/MLi-2), and with Rebastinib (RCKW/Rebastinib). The LRRK2^RCKW^ structure was built based on the reported cryo-EM structure (pdb:6VNO), and binding of the inhibitors was modeled by superimposing the reported respective kinase/inhibitor structures [12,31]. Ten replicated simulations were carried out for each of the models (RCKW, RCKW/MLi-2, and RCKW/Rebastinib). Both inhibitors stayed in the binding pocket throughout the simulation, and the RMSD value of the inhibitors indicated that both inhibitors bound to the LRRK2 protein stably. MLi-2, which is an ATP analog, occupied the ATP binding pocket of the LRRK2 kinase domain and was capped by the Gly-rich loop (aa 1886-1893) (Fig 3A). The morpholine group in MLi-2 was surrounded by residues in the hinge/linker region (aa 1949-1954), and R1895 on the β2 strand. L2001 and L1985, two C-spine residues, clamped to MLi-2 (Fig 3B) and E1948, at the end of β5, formed a stable H-bond with MLi-2. A1950 also interacted with MLi-2 through an H-bond to its backbone in some of the frames. Binding of MLi-2 stabilized the DYG-in/BLBplus conformation [32]. The LRRK2 kinase was in a DYG-In conformation for ninety percent of the simulation time (Fig S1).

**Fig 3.**
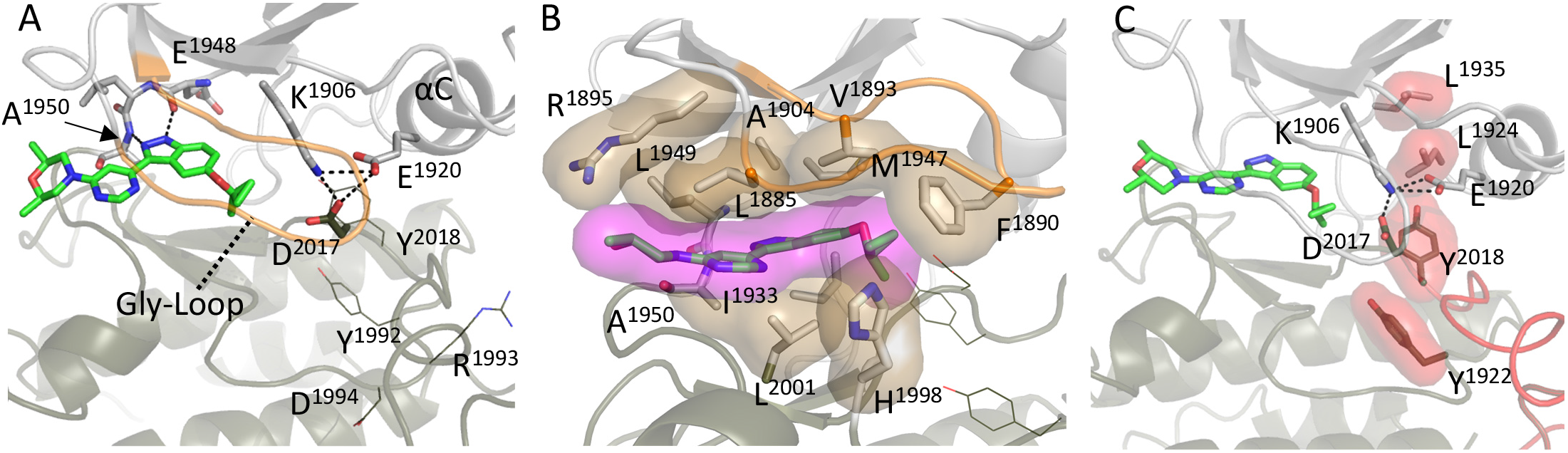
Structure of the binding pocket of the LRRK2 kinase domain with type I inhibitor. (A) MLi-2 binds in the ATP binding pocket of the kinase domain and forms hydrogen bonds with residues R1895 and E1948. (B) Hydrophobic interactions occur between MLi-2 and residues in the Gly-rich loops and the hinge region are highlighted. Two hydrophobic residues in particular, Phe1890 in the G-Loop and Leu2001, Phe2003 of the C-lobe fuse the N- and C-Lobes together. (C) The MLi-2 bond LRRK2 kinase samples the active-like conformation where the regulatory triad (K1906, E1920 and D2017) and regulatory spine (L1935, L1924, Y2018 and Y1922) are assembled. Structures of inhibitors are based on models of 5OPB.

The binding of MLi-2 also promoted the assembly of the highly conserved regulatory triad, the salt bridges that define every active kinase (Fig 3A) [33]. The salt bridges between two conserved residues in the N-Lobe, E1920 in the αC helix and K1906 in β3, and D2017 in the DYG motif in the C-lobe are essential for active forms of kinases. The free energy profiles along the distances of the two salt bridges are shown in Fig 4. Compared with the WT system (Fig 4A), the triad is more stable in the MLi-2-bound system (Fig 4B), which does not form metastable states at a larger K1906-E1920 distance (5∼6 Å). This triad also helps to drive the assembly of the R-spine, another hallmark feature of active kinases (Fig 3C, supplementary movie 1) [34]. Consistent with HDX-MD data, the YRD motif, where D1992 is an R-spine residue, is also stabilized by MLi-2, shown by the smaller RMSD values of residues in the region of the YRD motif (Fig S2). In addition, MLi-2 stabilized residues in the hinge region. Another hallmark signature of an active kinase that is in a fully closed conformation is the position of the hydrophobic residue in the Glycine-rich Loop, which is F1890 in LRRK2. As seen in figure 3B, F1890 further stabilizes the hydrophobic bridge between the N- and C-Lobes.

**Fig 4.**
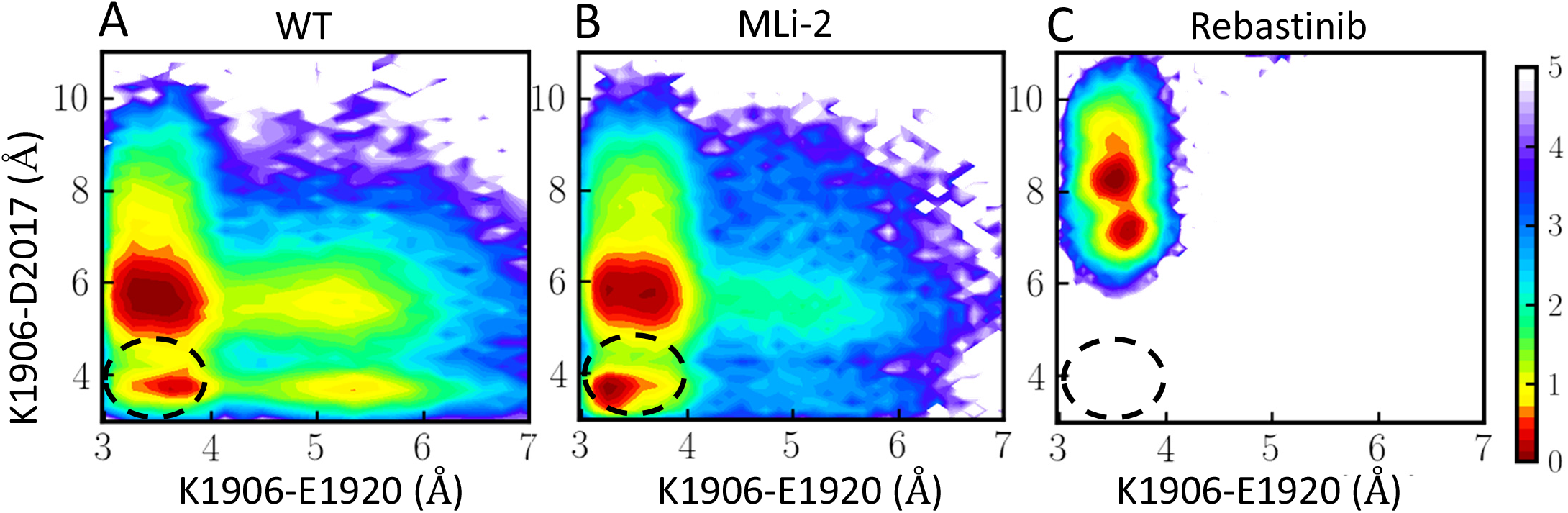
The formation of the regulatory triad. The two dimensional free energy profiles projected along two distance coordinates (A) LRRK2^RCKW^; (B) Mli-2 bound LRRK2^RCKW^; (C) Rebastinib bound LRRK2^RCKW^. The in x-axis shows the distance of K1906-E1920 and the y-axis shows the distance of K1906-D2017. Black circle indicates when the regulatory triad E1920-K1906-D2017 is formed. Binding of MLi-2 promote the assemble of regulatory triad while binding of Rebastinib inhibits it.

In contrast to MLi-2, Rebastinib is bound in the hydrophobic pocket adjacent to the ATP binding site, occupying the space that was occupied by Y2018 of the DYG motif in the MLi-2 structure (Fig 5A, 5B). This forces the LRRK2 kinase domain into the DYG-Out/BBAminus conformation [32]. The DYG motif was unable to flip back into the DYG-In orientation for the entire simulation time (Fig S1). Rebastinib breaks the hydrophobic bridge between the N- and C-Lobes, which defines the active kinase; it makes the closing of the active site cleft impossible. When they are uncoupled from the N-lobe, the dynamics of the DYG motif and the YRD motif were both also increased compared to the LRRK2^RCKW^ alone, as indicated by the higher RMSD values of residues in both motifs (Figure S2). The isoquinoline of the Rebastinib was positioned against the αC helix and pushed the αC helix further out. The E1920 on the αC helix is bound to Rebastinib through H-bonding with the two nitrogens on Rebastinib leaving it far away from D2017 in the DYG motif of the C-lobe. Although the salt bridge between K1906 and E1920, was stably present throughout the MD simulations, the regulatory triad never assembled (Fig 4C, supplementary movie 2).

**Fig 5.**
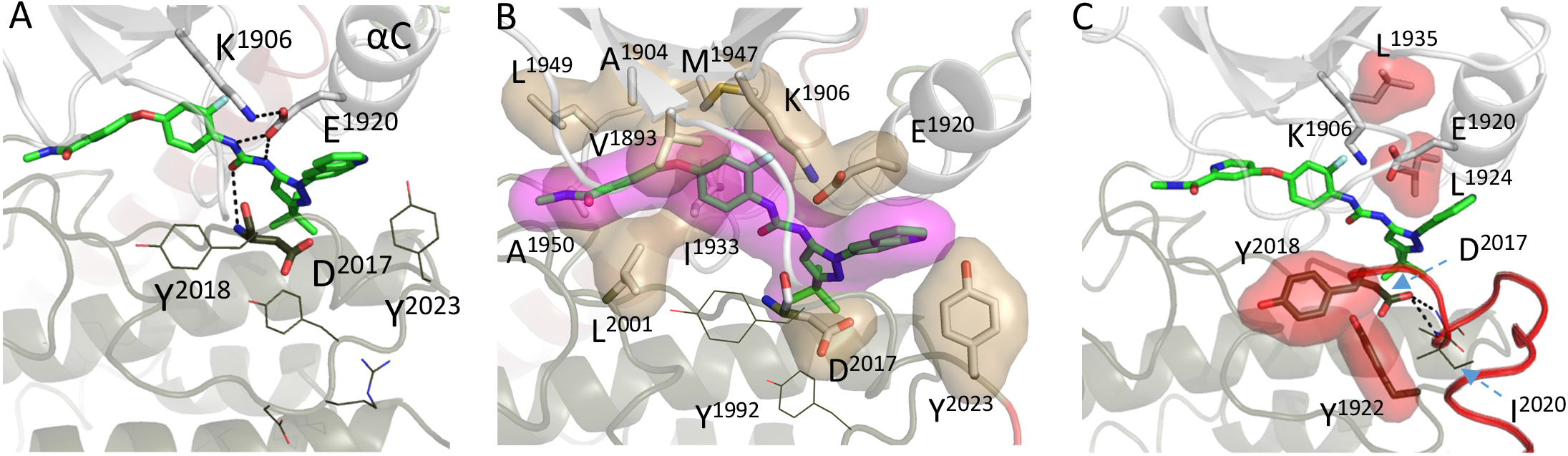
Structure of the binding pocket of the LRRK2 kinase domain with type II inhibitor. (A) Rebastinib displaces the Tyr of the DYG-motif and binds to residues E1920 and D2017. In addition, binding of Rebastinib keeps the αC helix in an extended out position. (B) Hydrophobic interactions with Rebastinib create a wedge between the N- and C-Lobes. (C) Rebastinib block the assembly of R-spine and the kinase domain is lock in the DYG-out, inactive conformation. The structures of inhibitors are based on models of 6MWE.

To further deduce how binding to the kinase inhibitors changes the conformation of LRRK2^RCKW^, we performed clustering analysis to extract representative structures from our simulation data. The first class of each condition, RCKW, RCKW/MLi-2, and RCKW/Rebastinib, are aligned by the stable helices, helix E and F, in the C-lobe of the kinase domain (Figure 6A). While the C-lobe aligns well except for the loop regions, the orientations of the N-lobe of the kinase domain are different when binding to the two inhibitors. This observation suggests that the N-lobe and the C-lobe move as independent rigid bodies while the WD40 domain remains stably anchored to the C-lobe in all of the structures solved so far (Figure S3). The Gly-rich loop and the αC Helix are closer to the C-lobe when LRRK2^RCKW^ binds to MLi-2 compared to the RCKW or RCKW /Rebastinib systems, showing a closed conformation.

**Fig 6.**
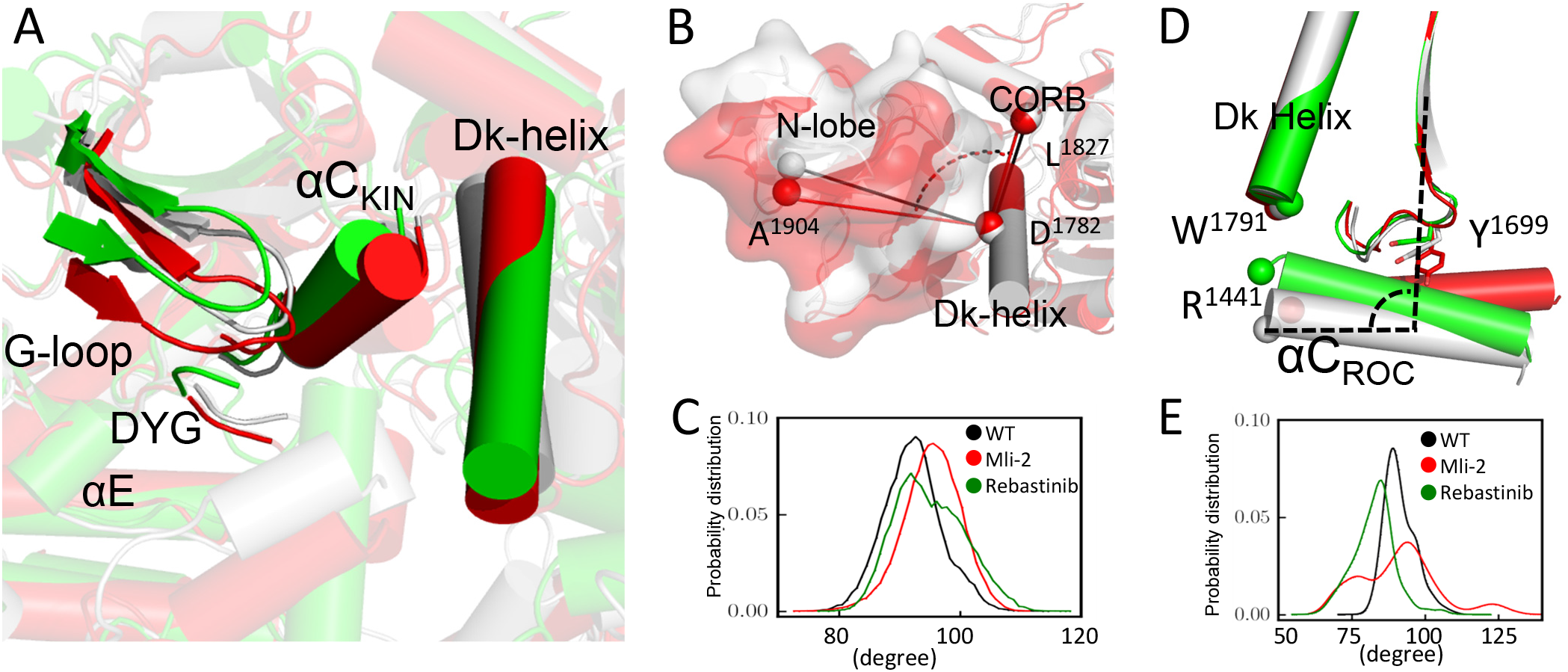
Clustering analysis of the MD conformations. (A) Representative structures from the first class of the clustering analysis are colored as follows: RCKW/WT (gray); RCKW/MLi2 (Red) ; RCKW/Rebastinib (green). Structures aligned by the αE and αF helices of the kinase domain show that the G-loop, αC and Dk-helix of RCKW/MLi-2 are closer to the C-lobe in the MLi-2-bound structure. In contrast, the RCKW/Rebastinib complex resembles the open conformation of the kinase domain. (B) The representative structures aligned by Dk-helix show that when MLi-2 is bound to RCKW, the top of the N-lobe moves away from the CORB domain compared to RCKW/WT. (C) The angle of A1904-D1782-L1827 was measured to show the relative orientation of the N-lobe toward the CORB domain. (D) The αC helix in the ROC-domain, which hinges at Y1669 (24), tilts towards the C-terminal end of the Dk-helix, when Rebastinib is bound. This tilting motion of RCKW/Rebastinib moves R1441 closer to W1791, whereas The R1441 moves way from the W1791 when bound to MLi-2. (E) The angle of R1693-W1434-R1441 was measured to show the tilting of the αC helix.

We note that the RCKW/MLi-2 structure is still distinctly different from the ATP-bound, fully closed, active conformation, based on PKA [35], where the Gly-loop folds over the nucleotide and the αC Helix is closer to the β-sheet core of the N-lobe (Fig S4). The glycine-loop including the Phe residue at the tip of the G-Loop folds over the MLi-2 while the Phe in the ATP:PKI bound structure of PKA (Phe54) folds over onto the DFG+1 residue and reinforces the hydrophobic latch between the N- and C-lobes. The tip of the αC helix, however, remains anchored to the Dk-helix although it is closer to the Activation Loop of the kinase domain. When binding to Rebastinib, the N-lobe is further away from the C-lobe compared to the MLi-2 bound conformation. By occupying the space that was filled by Y2018 in the MLi2-bound structure, Rebastinib essentially breaks the R-spine, which severs the N- and C-lobes (Fig 5C). In the inactive full-length LRRK2 structure, the DYG motif forms an inhibitory helix that prevents the assembly of R-spine (Fig S5). This inhibited structure, however, is distinct from the RCKW/Rebastinib structure; it does not correspond to the stable inhibited conformation that is found in the full length LRRK2.

### Inter-domain communications across LRRK2 are affected differently by the two inhibitors

Inter-domain interactions between the COR-B domain and the kinase domain are known to be involved in regulating the activation of LRRK2 [30]. When the kinase domain is stabilized in the closed conformation by MLi-2, the N-lobe of RCKW/MLi-2 rotates along the Dk-helix of the COR-B domain and moves closer to the COR-B domain (Figure 6B and 6C). More interactions can be identified between the Dk-helix and the COR-B loop (residues 1721–1725) (Fig S6A). In agreement, the COR-B loop also shows smaller deuterium uptake when LRRK2^RCKW^ binds to MLi-2 (Fig S6B). This orientation resembles the active LRRK2 structure recently solved by Sun’s group [27]. In contrast to MLi-2, binding of the Rebastinib did not alter the N-lobe orientation, and the HDX-MS result also shows no protection effect on the COR-B loop. We also analyzed the reported “seesaw-like” motion of the ROC αC helix relative to the COR-B domain, which was shown to be related to LRRK2 activation [27]. In the inactive conformation, the C-terminal end of the ROC αC helix stays closer to the C-terminal end of the Dk-helix. R1441 at the C-terminal end of the ROC αC helix, in particular, can bind to W1791 at the C-terminus of the Dk-helix anchoring the ROC αC helix to the surface of the COR-B domain in the inactive state. Rebastinib binding stabilizes this inactive conformation and tilts the ROC αC helix in the direction towards the Dk-helix (Fig 6D, 6E). On the other hand, binding to MLi-2 stabilizes an active-like conformation, where the C-terminal end of the ROC αC helix moves away from the Dk-helix. In this way MLi-2 prevents the interaction between R1441 and W1791 thus disrupting the anchoring of the ROC αC helix to the Dk-helix. These changes also most likely promote interactions with the Activation Loop of the kinase domain, which is highly dynamic and lies in close proximity to this region.

The dynamic changes related to the domains that flank the kinase domain was also captured by HDX-MS (Fig 7). The peptide that covers the C-terminus of the ROC αC helix (residues 1436-1449) shows increased deuterium uptake in the presence of MLi-2. This is consistent with the MD results showing that the interaction between W1791 and R1441 was disrupted by the binding of MLi-2, releasing the ROC αC helix and making it more dynamic and solvent accessible. Interestingly, two PD mutations are included in this important peptide, R1441 and N1437. In addition, the decreased deuterium uptake of the C-terminus of the Dk-helix (residues 1788-1795) can be attributed to its increased interactions with the Activation Loop observed in the simulation that shielded these residues from the solvent (Fig S7). Binding of the Rebastinib does not show any effect on the deuterium uptake of either the ROC αC helix or the Dk-helix. However, peptides that cover the residues at the interface of ROC-CORA (residues 1391-1401), CORA-CORB (residues 1666-1673) and ROC-CORB (residues 1467-1484) show decreased deuterium uptake in the presence of Rebastinib, while binding to MLi-2 has no protection effect. This suggests that the anchoring of ROC αC helix to the Dk-helix, not only influences the communication that takes place between the kinase domain and the CORB domain, but also alters the cross-talk within the ROC, CORA, and CORB domains.

**Fig 7.**
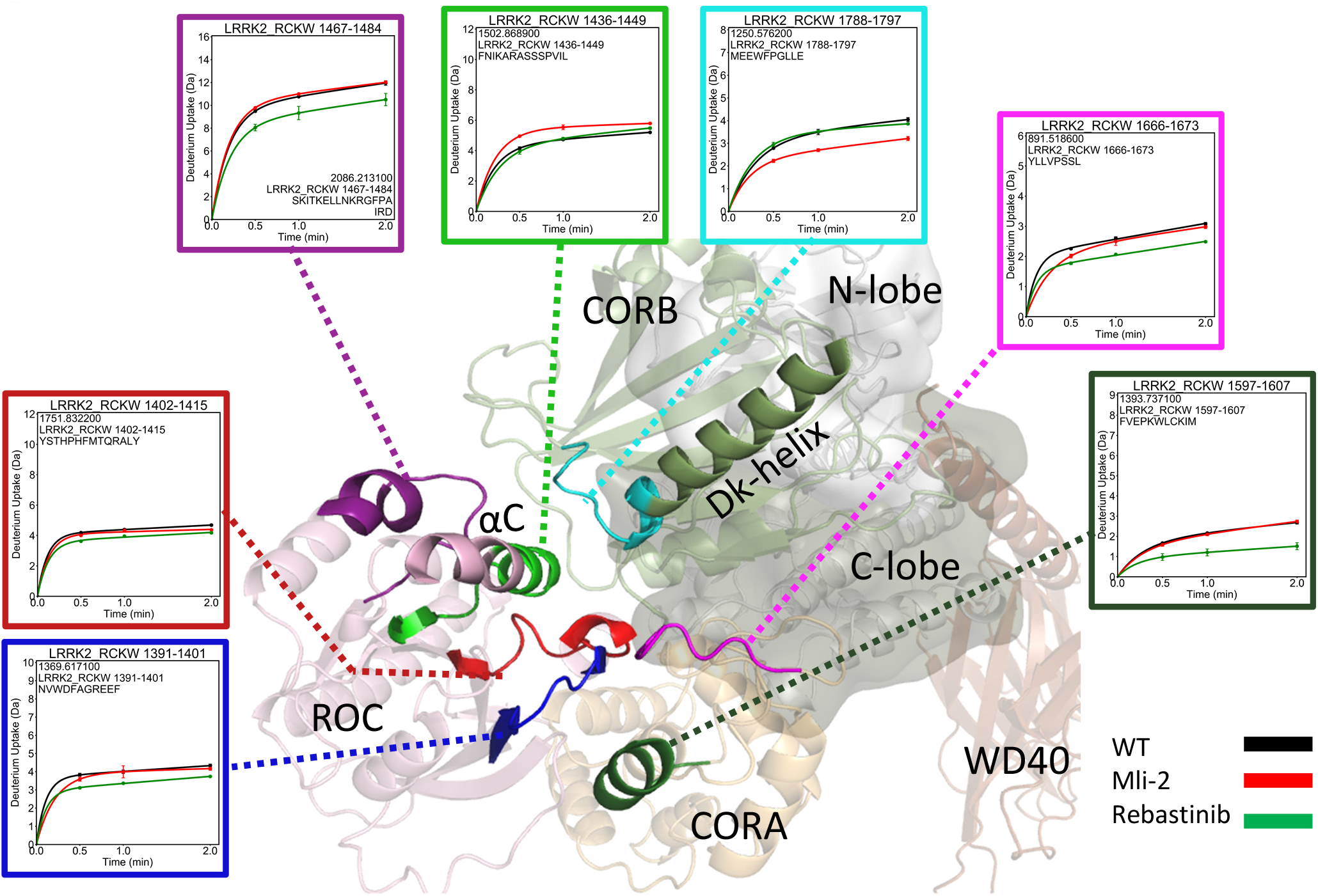
Deuterium uptake of LRRK2 at the ROC, CORA, and CORB interfaces. The deuterium uptake of selected peptides is plotted and mapped on the LRRK2^RCKW^ model. When binding to MLi-2, the peptide that includes the ROC αC helix shows increased its deuterium uptake while uptake is decreased in the peptide from the Dk-helix. For peptides at the domain interfaces, binding of Rebastinib globally reduces their deuterium uptake while MLi-2 binding has no effect.

### Type I and Type II inhibitors shift the energy landscapes for the LRRK2 conformational space differently

Using GaMD simulations we had previously captured the breathing dynamics of LRRK2^RCKW^ where the ROC-CORA, CORB, N-lobe and C-lobe-WD40 domain move as rigid bodies. The simulations showed how these rigid bodies fluctuate to create extended and compact conformations. The relative movement that closes the cleft shaped by the CORA, CORB and kinase domains upon activation was also captured by the cryo-EM structure [27]. To explore how binding of MLi-2 and Rebastinib affect the breathing dynamics of LRRK2_RCKW_ as well as the kinase domain equilibrium, we computed a 2-dimensional energy landscape for each simulation condition (Fig 8 A-C). The open and closed conformations of the kinase domain were measured by the relative position of the N- and C-lobes, which correlate with the inactive conformation where the N- and C-Lobes are uncoupled and the active-like conformation of the kinase domain where the two lobes come together (Fig 8D). For all of the simulation conditions, the kinase domain of LRRK2^RCKW^ toggles between open and closed conformations (y-axis of Figure 8A-C). The MLi-2-bound LRRK2^RCKW^ has an energy minimum at 28.5 Å, smaller than the minimum positions of the WT LRRK2^RCKW^ (29.2 Å) and Rebastinib-bound LRRK2 ^RCKW^ (29.8 Å).

**Fig 8.**
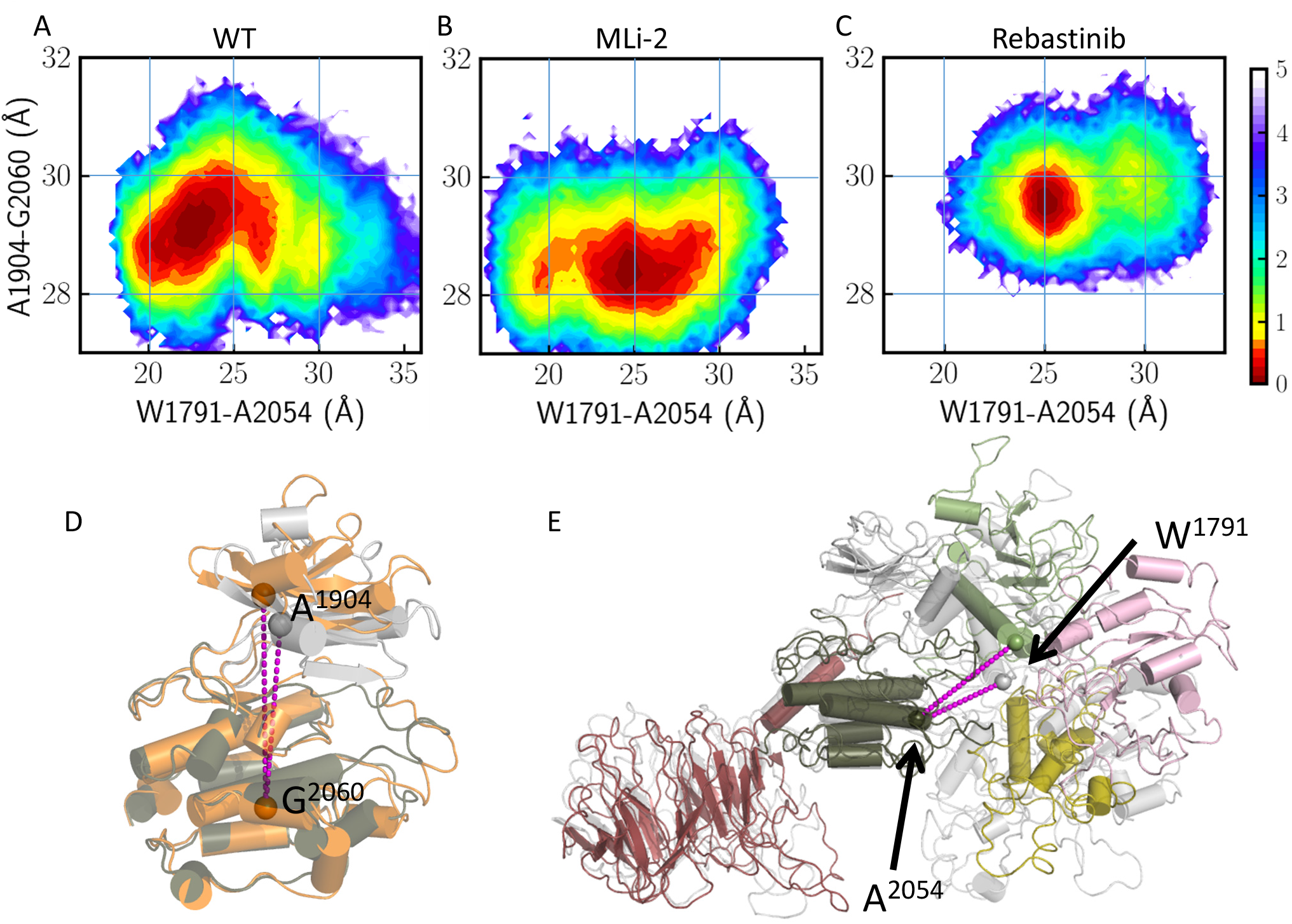
Binding of the inhibitors alter LRRK2 dynamics. The conformational free-energy landscape of RCKW/WT(A), RCKW/MLi-2 (B) and RCKW/Rebastinib (C). The distance from the N-lobe (A1904) to the C-lobe (G2060) represents the open (orange) and closed (gray) conformations of the kinase domain (D), while the distance between CORB (W1791) and C-lobe (A2054) represents the extended (colored) or compact (gray) conformation of LRRK2^RCKW^ (E). Both closed/compact and open/extended states are sampled by RCKW/WT. Binding of MLi-2 stabilizes the kinase domain in a closed closed/compact state. In contrast, the Rebastinib-bound LRRK2^RCKW^ is trapped in an open/extended state and is less dynamic.

The distance between the C-lobe and the CORB domain (the distance between A2054 and W1791 in Figure 8E) was measured to demonstrate the large-scale breathing motion. Both the extended and compact conformations of LRRK2_RCKW_ were sampled according to the distributions along the x-axis of Figure 8A-C. MLi-2-bound LRRK2 exhibits two local energy minima, one at 20 Å, representing a more compact configuration than that at the WT minimum (23 Å). The distribution along the x-axis for Rebastinib-bound LRRK2 shifts to a more extended configuration and has a minimum at 25 Å. We also calculated the 2D energy profiles using the C-lobe - CORA distance and the kinase conformational coordinate in Figure S5. Similarly, only MLi-2-bound LRRK2^RCKW^ has a population representing a very compact LRRK2 while the Rebastinib-bound LRRK2^RCKW^ shows minimal population for the compact configuration.

Based on the three energy profiles, the closed, active-like conformation of the kinase domain is in concert with the compact conformation of LRRK2^RCKW^ and vice versa, illustrating the correlation between the dynamics of the kinase domain and the breathing dynamics of LRRK2^RCKW^. Binding of MLi-2 shifts the kinase domain into a closed/compact state, and disrupts the coherent movement between the closed/compact and open/extended states (Fig 8B). MLi-2 bound LRRK2^RCKW^ was able to sample smaller distance distribution between the C-lobe and COR-B. On the other hand, binding of Rebastinib locks the kinase domain in an open, inactive state, which corresponds to the extended state, and the compact RCKW conformation is rarely sampled (Fig 8C and S8).

## Discussion

The function of LRRK2 is mediated by two finely tuned regulatory switches, a kinase domain and a GTPase (ROC) domain, which regulate how LRRK2 toggles between its active and inactive states [30,36-38]. The active and inactive states of LRRK2 not only correlate with its kinase activity, but also reflect the conformational changes that are involved in mediating the oligomeric state of LRRK2 as well as its interactions with other proteins where it functions as a scaffold that mediates a variety of multi-protein complexes [39]. We have shown that the kinase domain serves as a central hub for inter-domain communication and is the major driver for LRRK2’s conformational transitions [29,40]. Previous studies showed that different distributions of LRRK2 in the cell and different phenotypes are captured in the presence of type I or type II kinase inhibitors [20,21]. The study of Rab protein phosphorylation with type I and type II inhibitors also indicates that the function of LRRK2 is more than the activity of the kinase domain. The conformation status of LRRK2 that is mediated by the kinase domain also plays a crucial role in LRRK2 function[23]. In this study, we used GaMD simulations coupled with HDX-MS to analyze the dynamic changes of LRRK2 and its inter-domain allosteric communications when the kinase domain is locked into a closed, active-like conformation or an open, inactive conformation by binding of type I and type II inhibitors, respectively. Even though both MLi-2 (type I) and Rebastinib (type II) both inhibit the kinase activity with high affinity, they drive LRRK2 into different conformational states. Binding of MLi-2 reduced the solvent accessibility of the kinase domain not only in the active site cleft where the MLi-2 is directly docked, but also in regions in the C-lobe that lie far from the active site cleft but nevertheless interact with other domains. This can be attributed to the fact that the kinase domain assumes a stable, compact and more closed conformation in the presence of MLi-2. Although Rebastinib shows similar but more localized protection in the N-lobe of the kinase domain, the overall the deuterium uptake was increased in the C-lobe of the kinase domain. The HDX-MS data indicates that binding of Rebastinib traps the kinase domain in an open conformation, which separates the C-lobe of the kinase domain from the N-lobe making it more solvent exposed and more dynamic. The simulation also shows that the overall dynamics of the kinase domain is reduced by MLi-2 where the closed, active-like conformations are sampled more frequently. In contrast, binding of Rebastinib results in a more dynamic kinase domain that favors an open, inactive conformation where the N- and C-Lobes are uncoupled.

In addition to the kinase domain, the changes in conformation and dynamics that are caused by binding of kinase inhibitors extend beyond the kinase domain in LRRK2^RCKW^. In this study, we showed the distinct roles of the Dk-helix in the CORB domain, which is anchored to the αC helix in the N-Lobe of the kinase domain and bridges the allosteric conformational changes between the kinase and ROC domains. Our results are in agreement with the recent report showing that bridging of the ROC domain and the Activation Segment in the kinase domain is different in the active and inactive structures [27]. When the kinase is locked in an open conformation by Rebastinib, the communication between the N- and C-lobes of the kinase domain is disrupted, which is consistent with the N-Lobe functioning as an independent rigid body [26]. The C-lobe of the kinase becomes more dynamic and uncoupled from the CORB domain. Specifically, the C-terminal end of the Dk-helix in CORB is anchored to the αC helix of the ROC domain via direct interaction of W1791 with R1441. The αC helix in the COR domain is stabilized in a tilted direction, which pivots at residue Y1699. The interactions between the CORB-CORA, CORB-ROC, and the CORA-CORB domains are more stable based on the reduced solvent accessibility of the peptides that are located at these interfaces. LRRK2^RCKW^ is stabilized in an extended state by Rebastinib whereas, binding of MLi-2 stabilizes a closed, active-like conformation of the kinase domain. With MLi-2 the overall dynamic features of the kinase domain are reduced, and the disordered regions surrounding the Activation Segment become more ordered, and, as a consequence, more extensive interactions can be identified between the Dk-helix and the kinase domain. The compact conformation of LRRK2^RCKW^ is thus stabilized by MLi-2 (Figure 9). We demonstrate that the intrinsic interactions within and between domains lead to different dynamics and conformational states of LRRK2 that are driven by different kinase inhibitors. The intermediate states responsible for the allosteric regulation in the kinase and flanking domains can be new targets for therapeutic intervention. Our results also indicate that the inhibitors can be used to trap LRRK2^RCKW^ and potentially full length LRRK2 in specific states for future functional studies.

**Figure 9.**
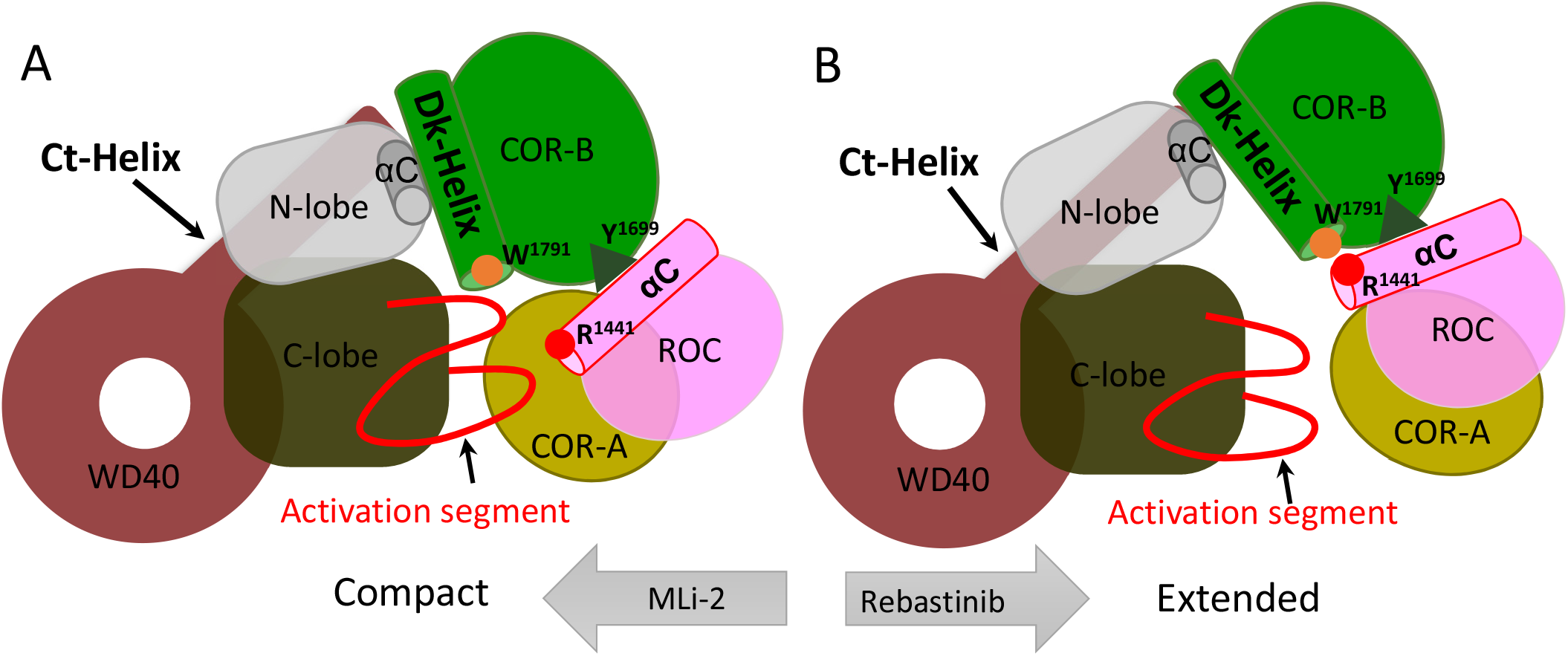
Cartoon representation of the compact and extended states of LRRK2^RCKW^. The kinase domain is stabilized in a closed conformation when the LRRK2^RCKW^ is in a compact conformation, where the ROC, CORA and CORB domains are closer to the C-lobe of the kinase domain. In this closed conformation the αC helix in the COR domain tilts away from the Dk-Helix. (B) The open, inactive conformation of the kinase domain promotes the extended conformation of LRRK2^RCKW^, and in this conformation the αC helix in the COR domain is closer to W1295 in the Dk-Helix of the CORB domain.

## Material and method

### Hydrogen-deuterium exchange mass spectrometry

LRRK2^RCKW^ proteins were expressed and purified from Sf9 cell [21]. Hydrogen/deuterium exchange mass spectrometry (HDX-MS) was performed using a Waters Synapt G2Si equipped with nanoACQUITY UPLC system with H/DX technology and a LEAP autosampler. The LRRK2_RCKW_ concentration was 5 μM in LRRK2 buffer containing: 20 mM HEPES/NaOH pH 7.4, 800 mM NaCl, 0.5 mM TCEP, 5% Glycerol, 2.5 mM MgCl_2_ and 20 μM GDP. The deuterium uptake was measured in LRRK2 buffer in the presence and absence of the kinase inhibitor MLi-2 (50 μM) or Rebastinib (50 μM). For each deuteration time, 4 μL complex was equilibrated to 25 °C for 5 min and then mixed with 56 μL D_2_O LRRK2 buffer for 0, 0.5, 1 or 2 min. The exchange was quenched with an equal volume of quench solution (3 M guanidine, 0.1% formic acid, pH 2.66). The quenched sample (50 μL) was injected into the sample loop, followed by digestion on an in-line pepsin column (immobilized pepsin, Pierce, Inc.) at 15 °C. The resulting peptides were captured on a BEH C18 Vanguard pre-column, separated by analytical chromatography (Acquity UPLC BEH C18, 1.7 μM, 1.0 × 50 mm, Waters Corporation) using a 7-85% acetonitrile gradient in 0.1% formic acid over 7.5 min, and electrosprayed into the Waters SYNAPT G2Si quadrupole time-of-flight mass spectrometer. The mass spectrometer was set to collect data in the Mobility, ESI+ mode; mass acquisition range of 200–2,000 (m/z); scan time 0.4 s. Continuous lock mass correction was accomplished with infusion of leu-enkephalin (m/z = 556.277) every 30 s (mass accuracy of 1 ppm for calibration standard). For peptide identification, the mass spectrometer was set to collect data in MS^E^, ESI+ mode instead.

The peptides were identified from triplicate MS^E^ analyses of 10 μM LRRK2_RCKW_, and data were analyzed using PLGS 3.0 (Waters Corporation). Peptide masses were identified using a minimum number of 250 ion counts for low energy peptides and 50 ion counts for their fragment ions. The peptides identified in PLGS were then analyzed in DynamX 3.0 (Waters Corporation) using a cut-off score of 6.5, error tolerance of 5 ppm and requiring that the peptide be present in at least 2 of the 3 identification runs. The peptides reported on the coverage maps are those from which data were obtained. The relative deuterium uptake for each peptide was calculated by comparing the centroids of the mass envelopes of the deuterated samples vs. the undeuterated controls [41]. For all HDX-MS data, at least 2 biological replicates were analyzed each with 3 technical replicates. Data are represented as mean values +/- SEM of 3 technical replicates due to processing software limitations, however the LEAP robot provides highly reproducible data for biological replicates. The deuterium uptake was corrected for back-exchange using a global back exchange correction factor (typically 25%) determined from the average percent exchange measured in disordered termini of various proteins[42]. Deuterium uptake plots were generated in DECA (github.com/komiveslab/DECA) and the data are fitted with an exponential curve for ease of viewing[43].

### Gaussian accelerated Molecular Dynamics (GaMD) simulation

The LRRK2_RCKW_ model for simulations was prepared based on the reported LRRK2_RCKW_ structure (PDB: 6VP6). Modeller was used to model the missing loops [44]. The Protein Preparation Wizard was used to build missing sidechains and model charge states of ionizable residues at neutral pH. Hydrogens and counter ions were added and the models were solvated in a cubic box of TIP4P-EW water molecules [45] and 150 mM KCl with a 10 Å buffer in AMBER tools. AMBER16 was used for energy minimization, heating, and equilibration steps, using the CPU code for minimization and heating and GPU code for equilibration. Parameters from the Bryce AMBER parameter database were used for phosphoserine and phosphothreonine [46]. Systems were minimized by 1000 steps of hydrogen-only minimization, 2000 steps of solvent minimization, 2000 steps of ligand minimization, 2000 steps of side-chain minimization, and 5000 steps of all-atom minimization. Systems were heated from 0 K to 300 K linearly over 200 ps with 2 fs time-steps and 10.0 kcal/mol/Å position restraints on protein. Temperature was maintained by the Langevin thermostat. Constant pressure equilibration with an 8 Å non-bonded cut-off and particle mesh Ewald was performed for 300 ps with protein and peptide restraints, followed by 900 ps of unrestrained equilibration. Gaussian accelerated MD (GaMD) was used on GPU-enabled AMBER16 for enhance conformational sampling [47]. GaMD applies a Gaussian distributed boost energy to the potential energy surface to accelerate transitions between meta-stable states while allowing accurate reweighting. Both dihedral and total potential boosts were used simultaneously. Potential statistics were collected for 2 ns followed by 2 ns of GaMD during which boost parameters were updated. Each GaMD simulation was equilibrated for 10 ns. For each construct, 10 independent replicates of 210 ns of GaMD simulation were run in the NVT ensemble, for an aggregate of 2.1 μs of accelerated MD.

Free energy landscapes were projected along selected conformational coordinates (e.g. Fig. 4C and E, and Fig. 6 A-C). For a 2D space of interest, we constructed a 2D histogram with a total number of M bins. The weighted histogram at bin *m* can be determined by

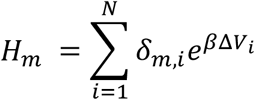

where *ΔV*_*i*_ is the boost potential at the *i*-th frame, *N* is the total number of frames, and *δ*_*m,i*_ is an indicator function that determines if frame *i* falls into bin *m*. The Maclaurin series expansion method

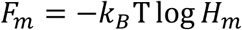

was used to approximate the exponential term [48]. The free energy profile can be determined by The “measure cluster” command in VMD was used to perform clustering analysis. The total number of clusters was 20. Molecular structures were rendered using PyMol.

## Acknowledgements

This work was supported by Michael J. Fox Foundation Grant 11425 (https://www.michaeljfox.org/) (to S.S.T.). The Synapt G2Si HD/X mass spectrometer was obtained from shared instrumentation NIH Grant S10 OD016234 (to S.S.). W.M and J.A.M were supported by National Institutes of Health grant R01-GM031749. The authors gladly acknowledge the GPU computing resources from the Triton Shared Computing Cluster at San Diego Supercomputer Center at UCSD. The funders had no role in study design, data collection and analysis, decision to publish, or preparation of the manuscript.

## Author Contributions

J.-H.W., W.M., and S.S.T. designed research; J.-H.W., S.S., and W.M. performed research; J.-H.W., W.M., S.S., J.W., and S.S.T. analyzed data; and J.-H.W., W.M., J.W., J.A.M. and S.S.T. wrote the paper. All author reviews the manuscript.

## Competing Interests statement

The authors have no competing interests.

## Supporting information

**S1 Fig. The conformation of DYG motif**. (A) The frequency of DYG-in or DYG-out conformation. Binding of Mli-2 promote the DYG-in conformation and binding of Rebastinib lock the DYG motif in a DYG-out conformation. (B) The DYG-in or DYG-out orientation is measured by the dihedral angle of O^A2016^-Cα^A2016^-Cα^Y2018^-Cγ^Y2018^.

**S2 Fig. The dynamic of kinase domain**. The RMSD distribution of selected regions (indicated in red) includes: hinge (residues 1948-1953), DYG motif (residues 2015-2021) and YRD motif (residues 1990-1995).

**S3 Fig. Clustering analysis of the MD conformations**. (A) Representative structures from the first class of the cluster analysis are colored as follows: RCKW/WT (gray); RCKW/MLi2 (Red); RCKW/Rebastinib (green). Structures aligned by the beta sheets of the N-lobe did not change much, except for small changes in the Gly-loop and the αC helix. The N-lobe moves as a rigid body during simulation.

**S4 Fig. Alignment of PKA and the reprehensive structure of LRRK2/MLi-2**. (A) The kinase domain of the representative structure of LRRK2^RCKW^/MLi-2 from the clustering analysis (red) aligned to the PKA catalytic subunit (pdb: 1atp) shown in gray. (B) Zoom in of the N-lobe of the kinase domain.

**S5 Fig. The regulatory spine of the Full-length LRRK2**. The R-spine is broken in the inactive full-length LRRK2 (pdb:7lhw). The inhibitory helix of the DFG motif prevent the assembly of R-spine.

**S6 Fig. Comparing the orientation of N-lobe and COR-B domain when LRRK2**^**RCKW**^ **binds to inhibitors**. (A) The N-lobe of the kinase domain moves closer to the COR-B loop (residues 1715-1732) and potentially increases the interactions among them. (B) The deuterium uptake of peptide 1715-1732 is reduced in the presence of MLi-2.

**S7 Fig. Interactions between kinase domain and CORB domain**. (A) Interactions between the activation loop (Q2022 and R2026) and Dk-helix (E1789) captured in the simulation. (B) Binding of MLi-2 promote the interactions of E1789-Q2022 and E1789-R2026.

**S8 Fig. Two dimensional (2D) free energy profiles projected along two distance coordinates** (A) LRRK2^RCKW^; (B) Mli-2 bound LRRK2^RCKW^; (C) Rebastinib bound LRRK2^RCKW^. The x-axis measures the distance between C-lobe (G2060) of the kinase domain and COR-A domain (N1577); The y-axis measures the distance between C-lobe (G2060) and N-lobe (A1904) of the kinase domain.

**S1 File. Simulation movie of MLi-2 bond LRRK2 kinase domain**.

**S2 File. Simulation movie of Rebastinib bond LRRK2 kinase domain**.

## Notes

### Competing Interest Statement

The authors have declared no competing interest.

