## supplemental figures for "Capturing differences in the regulation of LRRK2 dynamics and conformational states by small molecule kinase inhibitors"

Supplementary Figure 1

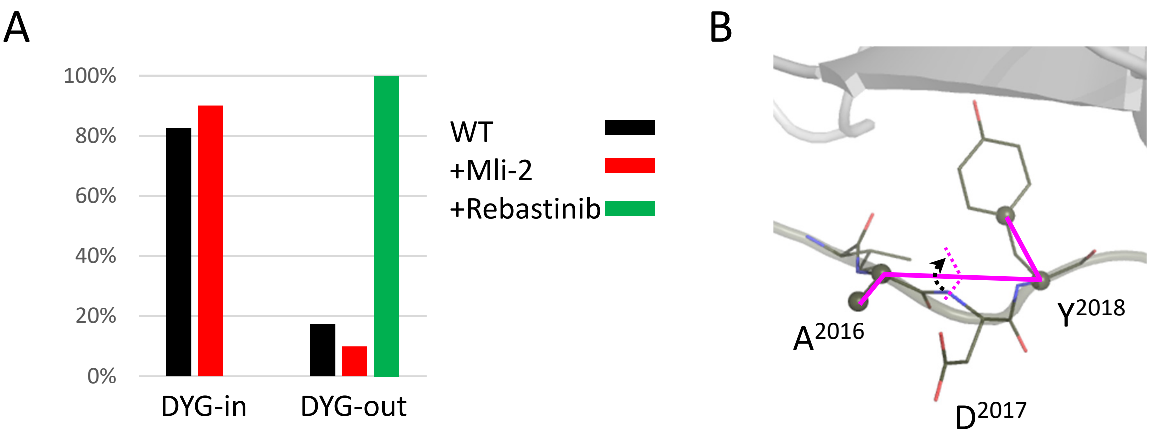


**Figure S1. The conformation of DYG motif**

(A) The frequency of DYG-in or DYG-out conformation. Binding of Mli-2 promote the DYG-in conformation and binding of Rebastinib lock the DYG motif in a DYG-out conformation. (B) The DYG-in or DYG-out orientation is measured by the dihedral angle of $O^{A2016}$-$C\alpha^{A2016}$-$C\alpha^{Y2018}$-$C\gamma^{Y2018}$.

Supplementary Figure 2

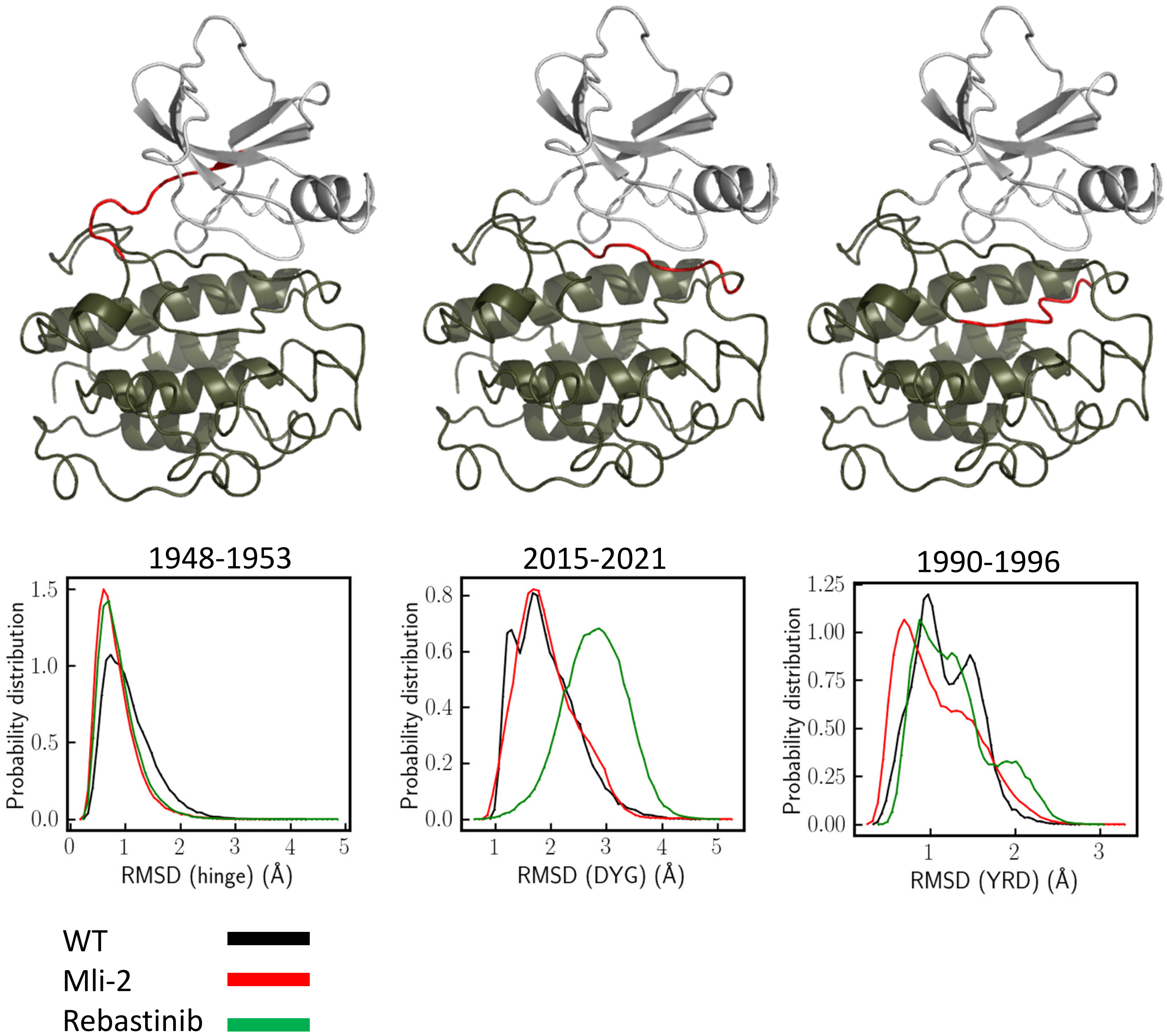


**Figure S2. The dynamic of kinase domain.** The RMSD distribution of selected regions (indicated in red) includes: hinge (residues 1948-1953), DYG motif (residues 2015-2021) and YRD motif (residues 1990-1995).

Supplementary Figure 3

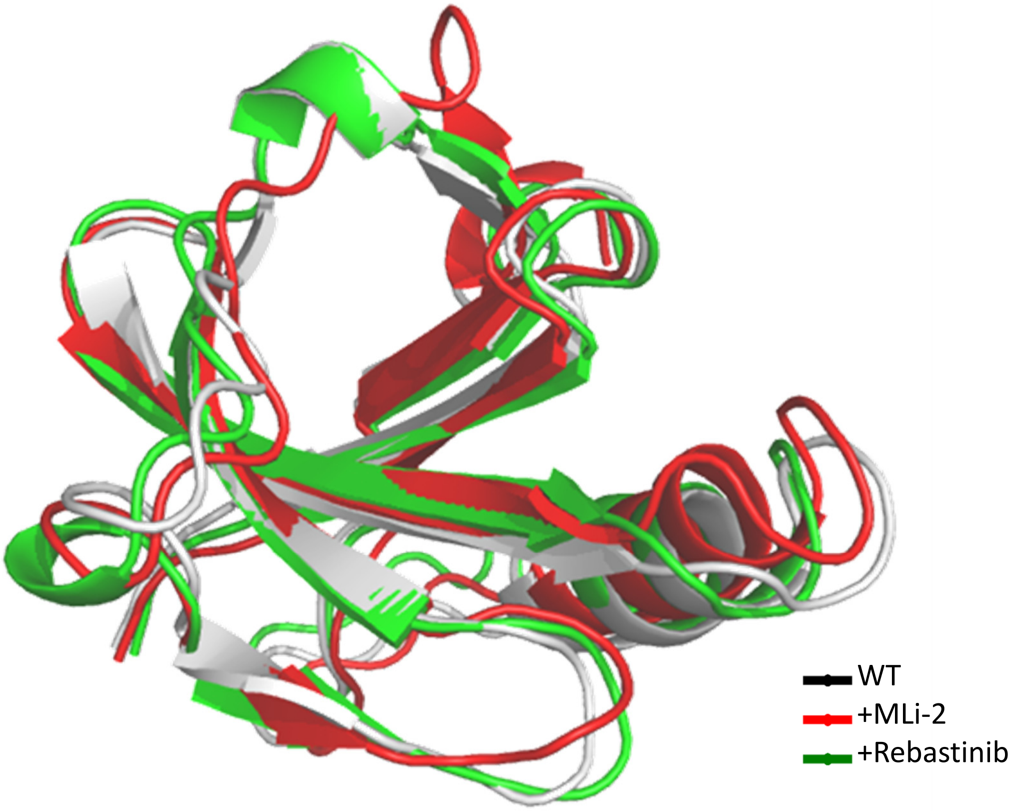


**Figure S3. Clustering analysis of the MD conformations.** (A) Representative structures from the first class of the cluster analysis are colored as follows: RCKW/WT (gray); RCKW/MLi2 (Red); RCKW/Rebastinib (green). Structures aligned by the beta sheets of the N-lobe did not change much, except for small changes in the Gly-loop and the αC helix. The N-lobe moves as a rigid body during simulation.

Supplementary Figure 4

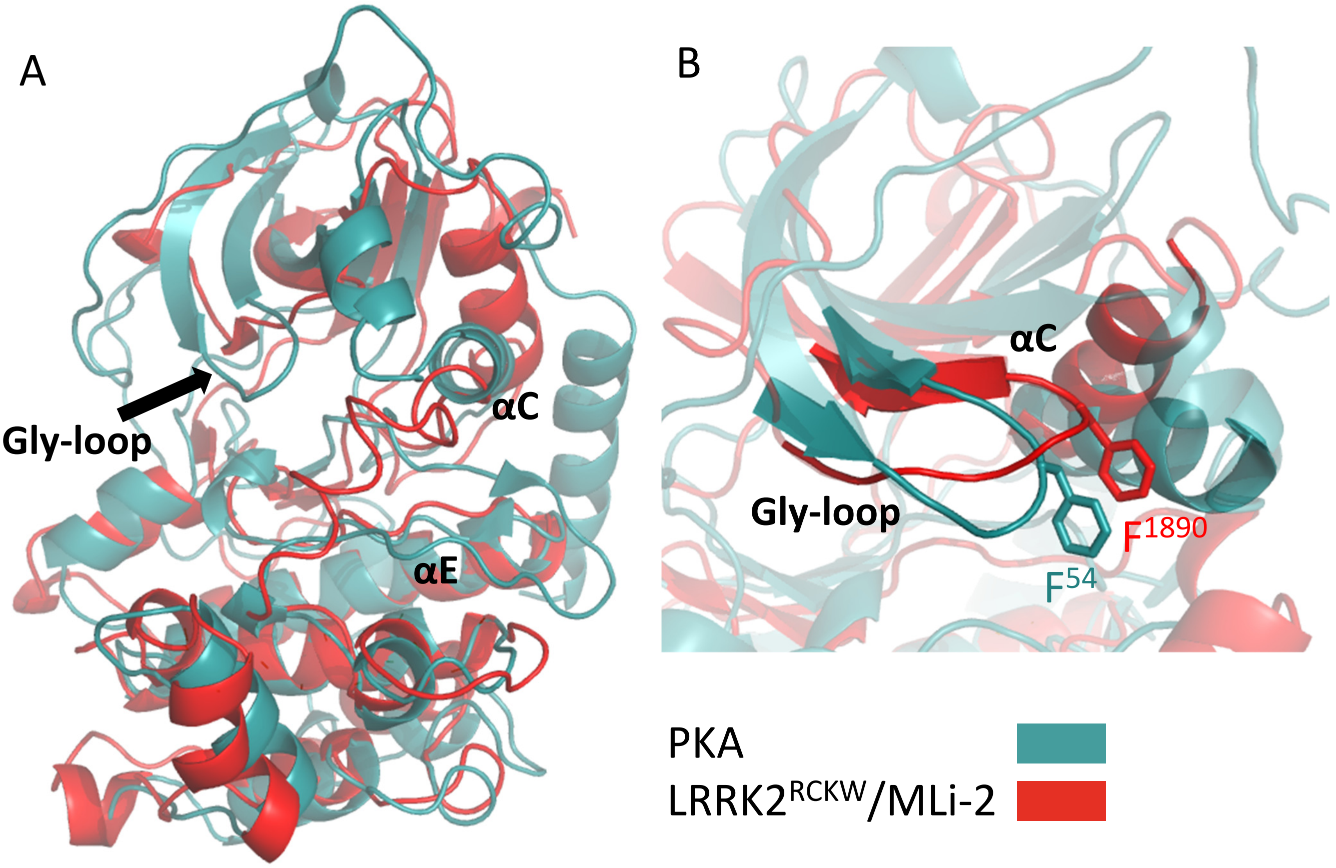


**Figure S4. Alignment of PKA and the reprehensive structure of LRRK2/MLi-2 .** (A) The kinase domain of the representative structure of LRRK2$^{RCKW}$/MLi-2 from the clustering analysis (red) aligned to the PKA catalytic subunit (pdb: 1atp) shown in gray. (B) Zoom in of the N-lobe of the kinase domain.

Supplementary Figure 5

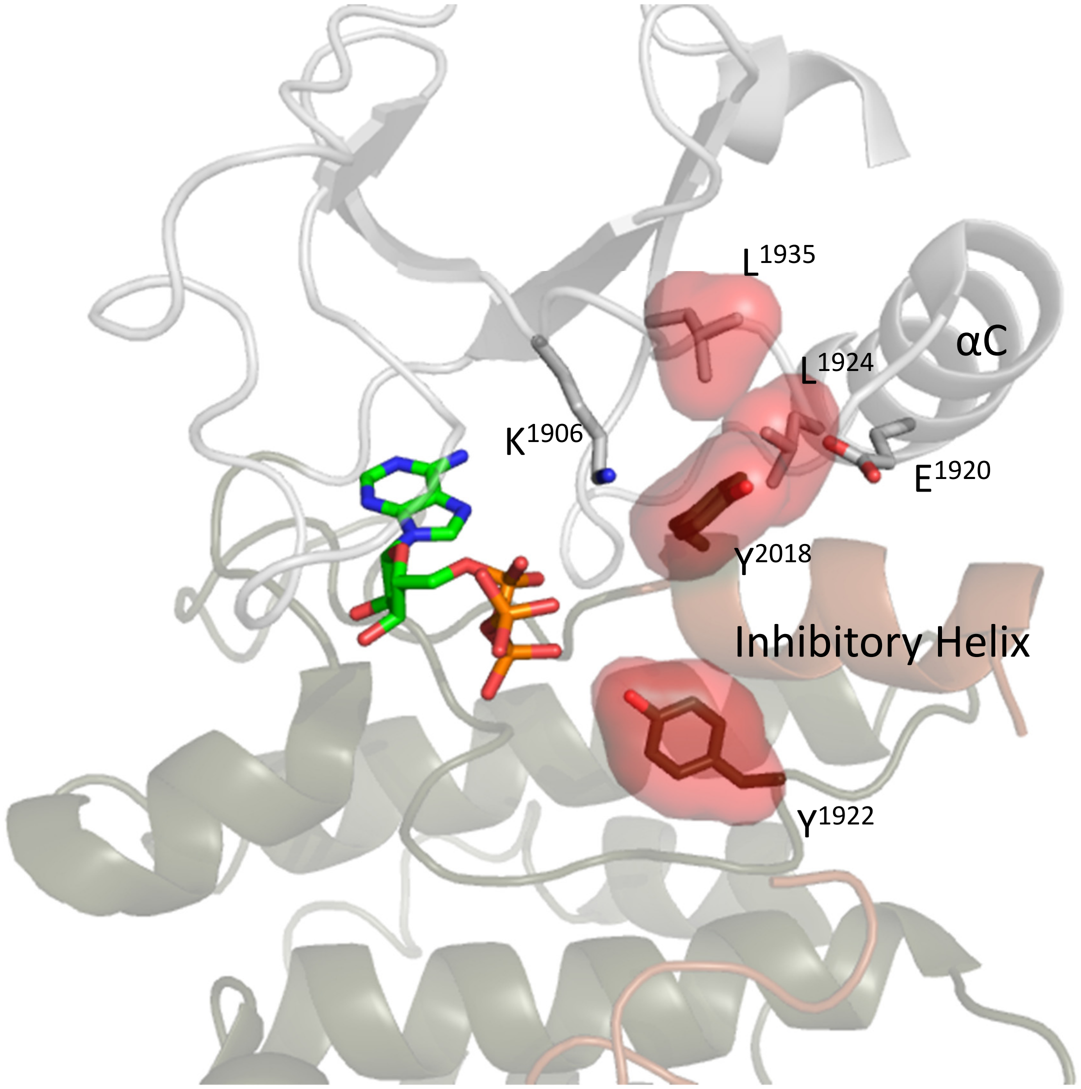


**Figure S5. The regulatory spine of the Full-length LRRK2.** The R-spine is broken in the inactive full-length LRRK2 (pdb:7lhw). The inhibitory helix of the DFG motif prevent the assembly of R-spine.

Supplementary Figure 6

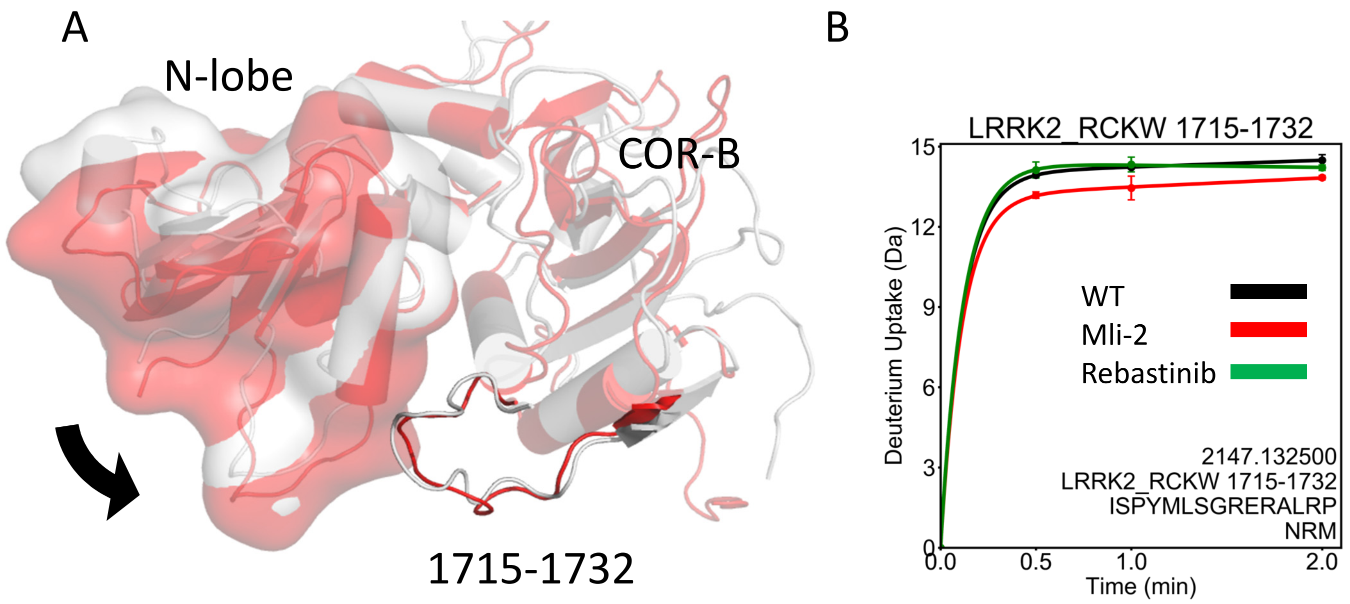


**Figure S6. comparing the orientation of N-lobe and COR-B domain when LRRK2$^{RCKW}$ binds to inhibitors.** (A) The N-lobe of the kinase domain moves closer to the COR-B loop (residues 1715-1732) and potentially increases the interactions among them. (B) The deuterium uptake of peptide 1715-1732 is reduced in the presence of MLi-2.

Supplementary Figure 7

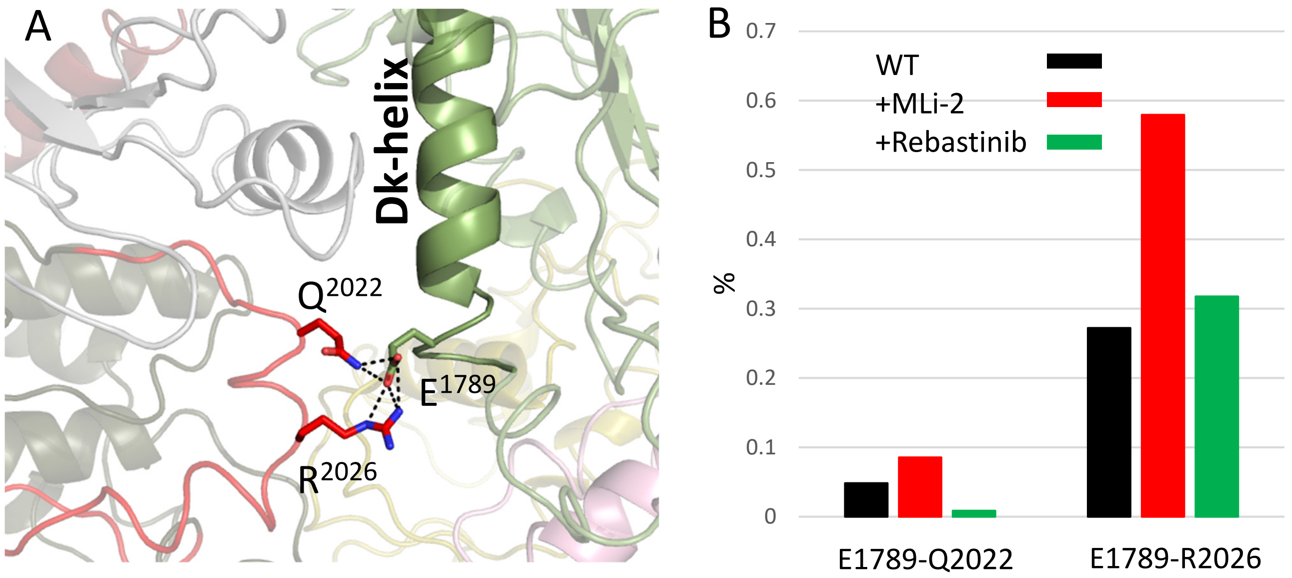


**Figure S7. Interactions between kinase domain and CORB domain.** (A) Interactions between the activation loop (Q2022 and R2026 ) and Dk-helix (E1789) captured in the simulation. (B) Binding of MLi-2 promote the interactions of E1789-Q2022 and E1789-R2026.

Supplementary Figure 8

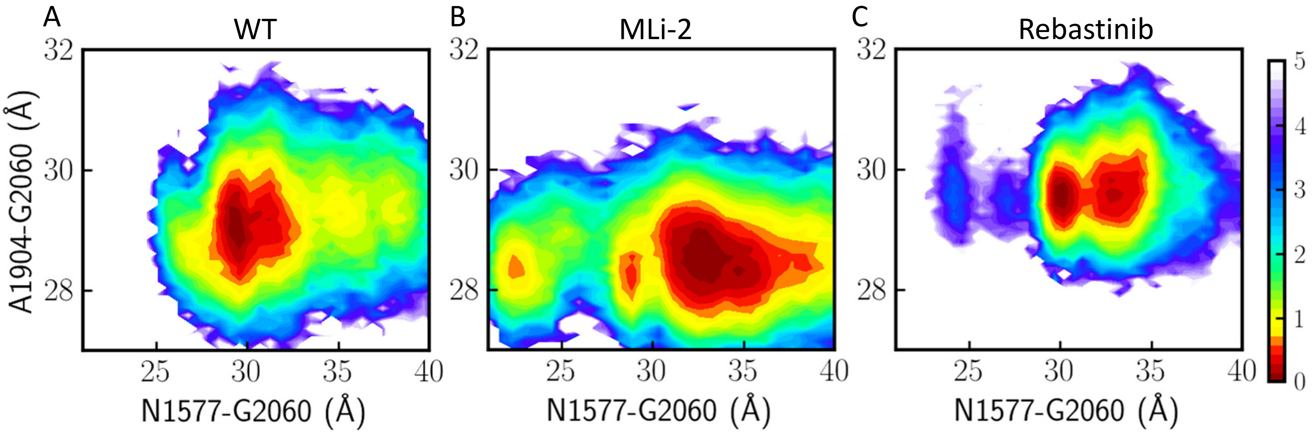


**Figure S8. Two dimensional (2D) free energy profiles projected along two distance coordinates** (A) LRRK2$^{RCKW}$; (B) Mli-2 bound LRRK2$^{RCKW}$; (C) Rebastinib bound LRRK2$^{RCKW}$. The x-axis measures the distance between C-lobe (G2060) of the kinase domain and COR-A domain (N1577); The y-axis measures the distance between C-lobe (G2060) and N-lobe (A1904) of the kinase domain.
